# Temporal distribution of deleterious variations influences the estimation of *F*_ST_

**DOI:** 10.1101/678011

**Authors:** Sankar Subramanian

## Abstract

Estimating the extent of genetic differentiation between populations is an important measure in population genetics, ecology and evolutionary biology. Fixation index or *F*_*ST*_ is an important measure, which is routinely used to quantify this. Previous studies have shown that *F*_*ST*_ estimated for selectively constrained regions was significantly lower than that estimated for neutral regions. By deriving the theoretical relationship between *F*_*ST*_ at neutral and constrained sites we show that an excess in the fraction of deleterious variations segregating within populations compared to that segregates between populations is the cause for the reduction in *F*_*ST*_ estimated at constrained sites. Using whole genome data, our results revealed that the magnitude of reduction in *F*_*ST*_ estimates obtained for selectively constrained regions was much higher for distantly related populations compared to those estimated for closely related pairs. For example, the reduction was 49% for comparison between European-African populations, 31% for European-Asian comparison, 16% for the Northern-Southern European pair and only 4% for the comparison involving two Southern European (Italian and Spanish) populations. Since deleterious variants are purged over time due to purifying selection, their contribution to the among population diversity at constrained sites decreases with the increase in the divergence between populations. However, within population diversity remain the same for all pairs compared above and therefore *F*_*ST*_ estimated at constrained sites for distantly related populations are much smaller than those estimated for closely related populations. Our results suggest that the level of population divergence should be considered when comparing constrained site *F*_*ST*_ estimates obtained for different pairs of populations.

## Introduction

Since the introduction of F-statistics by Sewall Wright (Wright 1951), fixation index or *F*_ST_ has been routinely used to measure the extent of differentiation between populations (Akey, et al. 2002; Beaumont and Balding 2004; Bersaglieri, et al. 2004; Barreiro, et al. 2008; Excoffier, et al. 2009; Keinan, et al. 2009; Xue, et al. 2009; Chen, et al. 2010; Wu and Zhang 2010; Sams and Hawks 2013; Vitti, et al. 2013). *F*_ST_ compares the heterozygosities within and between (or total) populations to measure the level of genetic structure among populations. Apart from being an integral part of the descriptive statistics to describe a population, *F*_ST_ has direct applications in conservation biology, ecology, evolutionary biology and clinical genetics. *F*_ST_ reveals the extent of genetic drift and the level of migrations between populations, which is useful to understand the population dynamics of an ecosystem (Whitlock and McCauley 1999). The level of differentiation in populations helps conservation biologists to measure risk of extinction of a population or species (Frankham, et al. 2002). Furthermore, *F*_ST_ is used to identify candidate genetic variants and genes associated with Mendelian and complex genetic diseases (Akey, et al. 2002; Barreiro, et al. 2008; Sams and Hawks 2013).

In evolutionary biology *F*_ST_ is used to detect the signature of positive selection (Beaumont and Balding 2004; Barreiro, et al. 2008; Excoffier, et al. 2009; Xue, et al. 2009; Bonhomme, et al. 2010; Chen, et al. 2010; Wu and Zhang 2010; Vitti, et al. 2013). Since adaptive mutations quickly spread through a population, their relative high frequency in one population (compared to the other) elevates the *F*_ST_ estimates. In whole genome analyses, *F*_ST_ is estimated using a sliding window to detect genomic regions showing high differentiation between populations (Vitti, et al. 2013). Although a number of previous studies have investigated the relationship between *F*_ST_ and positive selection only a handful of studies examined the influence of negative selection on *F*_ST_. A previous study reported lower *F*_ST_ for genic compared to nongenic SNPs (Barreiro, et al. 2008). The reduction in *F*_ST_ was more pronounced when only the amino acid changing nonsynonymous SNPs (nSNPs) were considered and a similar reduction was observed for mutations in disease-related genes. This suggests that purifying selection does not allow an increase in the frequency of SNPs, which could have led to the observed low *F*_ST_ (Nielsen 2005). Later a more systematic investigation was conducted to examine this issue using human genome data (Maruki, et al. 2012). This study grouped nSNPs based on the evolutionary rates of sites in which they were present and showed a positive correlation between the rates and *F*_ST_. Hence *F*_ST_ estimated for the nSNPs present in selectively constrained sites (with low rate of evolution) was much smaller than that estimated for those present in neutral sites with high evolutionary rates. A similar observation was made by another study on the populations of fruit flies from France and Rwanda (Jackson, et al. 2015). *F*_ST_ estimates obtained for long introns (known to be under high purifying selection) and conserved genes were typically lower than those estimated for short introns (under relaxed selective constraints) and less conserved genes.

Although the influence of purifying selection on *F*_ST_ estimates has been well documented, how exactly selective constraint affects *F*_ST_ estimations or the mechanism by which purifying selection influences these estimates is unclear. Furthermore, whether the magnitude of reduction in *F*_ST_ is depended on the divergence between populations or whether the magnitude of reduction is similar between closely related and distantly related populations is also unknown. To examine these, we first investigated the theoretical relationship between *F*_ST_ at neutral and constrained sites. Using the data from the 1000 genome project - Phase 3 (Auton, et al. 2015) we then estimated *F*_ST_ for pairs of populations with different levels of divergence such as Europeans-Africans, Europeans-Asians, Northern-Southern Europeans and two Southern European populations (Italians and Spanish).

## Materials and Methods

### Estimating the excess fraction of deleterious variants present within population

Heterozygosity in neutral and selectively constrained sites can be expressed as follows:

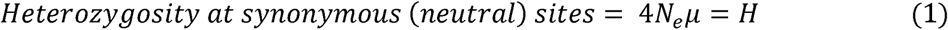

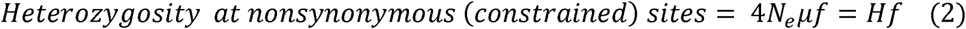

where *H, N*_*e*_ and *µ* are heterozygosity, effective population size and mutation rate respectively. The fraction of neutral and slightly deleterious mutations segregating in the population is denoted as *f*.

In terms of heterozygosity *F*_ST_ at synonymous sites (*F*_ST(S)_) can be expressed using Hudson et.al (Hudson, et al. 1992) as:

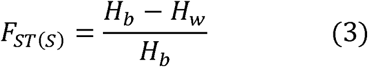

where *H*_*b*_ and *H*_*w*_ are synonymous site heterozygosity for between and within populations.

Using equation 2, *F*_ST_ at nonsynonymous sites (*F*_ST(N)_) is given as:

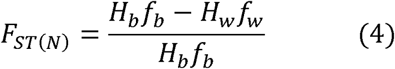

where *f*_*b*_ and *f*_*w*_ are fractions of neutral plus slightly deleterious nonsynonymous mutations segregating between and within populations respectively. For comparisons involving two populations *f*_*w*_ = (*f*_1_ + *f*_2_)/2 where *f*_*1*_ and *f*_*2*_ are neutral+deleterious fractions in populations 1 and 2 respectively. If *f*_*b*_ and *f*_*w*_ fractions of nonsynonymous mutations are equal, then we can show that *F*_ST_ at synonymous sites is equal to that at nonsynonymous sites as given below.

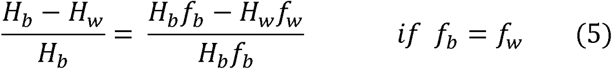

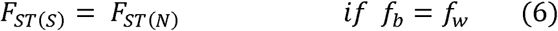

However, it is well known that the fraction of slightly deleterious mutations segregating within population is higher than that segregates between populations. This is because a much higher fraction of those segregating within population are young and yet to be purged from the population by natural selection. Therefore, *f*_*w*_ is expected to be higher than *f*_*b*_. Hence, we get,

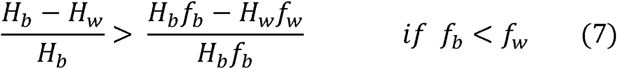

The above equation could be simplified by converting the fraction *f*_*w*_ in terms of *f*_*b*_ as:

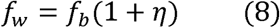

In the above equation *η* is the excess fraction of deleterious variations segregating within populations than that segregates between populations. Substituting this for *f*_*w*_we get,

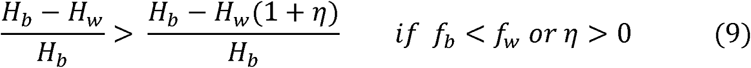

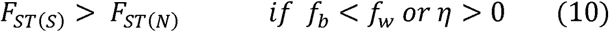

The above relationships clearly show that if *f*_*w*_ is higher than *f*_*b*_ or if there is an excess in the proportion of deleterious variations segregating within populations compared to that between populations (*η*) then *F*_ST_ of nonsynonymous sites will be smaller than that of synonymous sites. The magnitude of reduction in the *F*_ST_ of nonsynonymous sites could be quantified as:

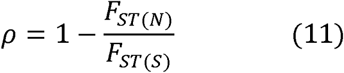

which is

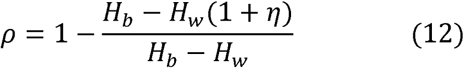

Equation 12 shows the theoretical relationship between *ρ* and *η*. However, using equation 11, *ρ* can be empirically estimated for the exome data using the estimates of *F*_ST_ at neutral 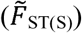 and constrained sites 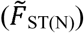.

A similar relationship for the Nei’s *F*_ST_ (*G*_ST_) for neutral and constrained sites is:

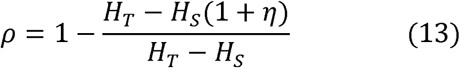

where *H*_*T*_ and *H*_*S*_ are heterozygosity for total and subpopulations.

### Population genome data

Whole genome data for 26 world-wide populations was downloaded from the 1000 genome project – Phase 3 (ftp://ftp.1000genomes.ebi.ac.uk/vol1/ftp/release/20130502/) (Auton, et al. 2015). Only biallelic single nucleotide polymorphisms (SNPs) from the autosomes were included for the analyses and the allele frequencies of each SNP in 26 populations were computed, which were used for estimating *F*_ST_ using the estimators described below. Pairwise FSTs were computed for the exomes of TSI (Italian)-YRI (Nigerian), TSI (Italian) – CHB (Chinese), TSI (Italian) – GBR (British) and TSI (Italian) – IBS (Spanish). To determine the magnitude of selective constraints at each site of the human genome we used a robust method, Combined Annotation-Dependent Depletion (CADD) that integrates many diverse annotations into a single measure (*C* score). (Kircher, et al. 2014). The precomputed C scores for each genome position are available at: http://cadd.gs.washington.edu/download/1000G_phase3_inclAnno.tsv.gz and these scores were mapped to the genotype data from the 1000 genome project. To identify derived alleles, orientations of SNVs were determined using the ancestral state of the nucleotides, which was inferred from six primate EPO alignments (Auton, et al. 2015).

### *F*_ST_ estimation

For estimating *F*_ST_ from human exome data we used two methods developed by Hudson et.al (Hudson, et al. 1992) and Nei (Nei 1973) and. We used the following estimators:

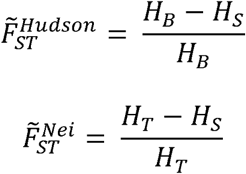

and

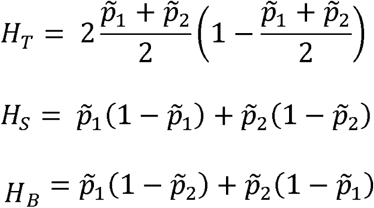

where *p*_1_, *p*_2_ are frequencies of the bialleles. To combine *F*_ST_ estimated for different SNPs of the genome we used the ratio of averages approach suggested by Bhatia et.al. (Bhatia, et al. 2013). To estimate the variance, we used a bootstrap resampling procedure with 1000 replicates.

## Results

### The effect of purifying selection on *F*_ST_

To examine the influence of purifying selection on *F*_ST_ we used European and African exome data from the 1000 genome project - Phase 3 (see methods). In order to examine the magnitude of selection pressure the Combined Annotation Dependent Depletion (CADD) score or *C*-score was used (Kircher, et al. 2014). Nonsynonymous SNPs were grouped into seven categories based on their *C*-scores. Figure 1A shows the relationship between selection pressure and *F*_ST_ estimated for synonymous (sSNPs) and nonsynonymous SNPs (nSNPs) using the exome data for the Italian (TSI) – Nigerian (YRI) pair. Clearly *F*_ST_ is the highest for the neutral sSNPs, which declines with increase in selection magnitude. *F*_ST_ estimate for highly constrained nSNPs with a *C*-score >30 was only 0.082, which is much smaller than that estimated for sSNPs (0.154). We introduced a measure, *ρ* to capture the magnitude of reduction in *F*_ST_ estimates of nSNPs compared to that of sSNPs (Equation 11 - see methods). Figure 1B that shows the positive relationship between the extent of selection constraint and magnitude of reduction of *F*_ST_ (*ρ*). The reduction in the *F*_ST_ estimate was only 2% for nSNPs under relaxed selection pressure (*C*-Score <=5), which increases with the magnitude of selection pressure. For highly constrained nSNPs (*C*-score >30), the reduction in *F*_ST_ was 47%, which is 24 times higher than that observed for nSNPs under relaxed constraint. Please note that the results shown were based on Hudson’s estimator and the results obtained using Nei’s estimator are given in the Supplementary information (Figures S1-S3).

**Figure 1.**
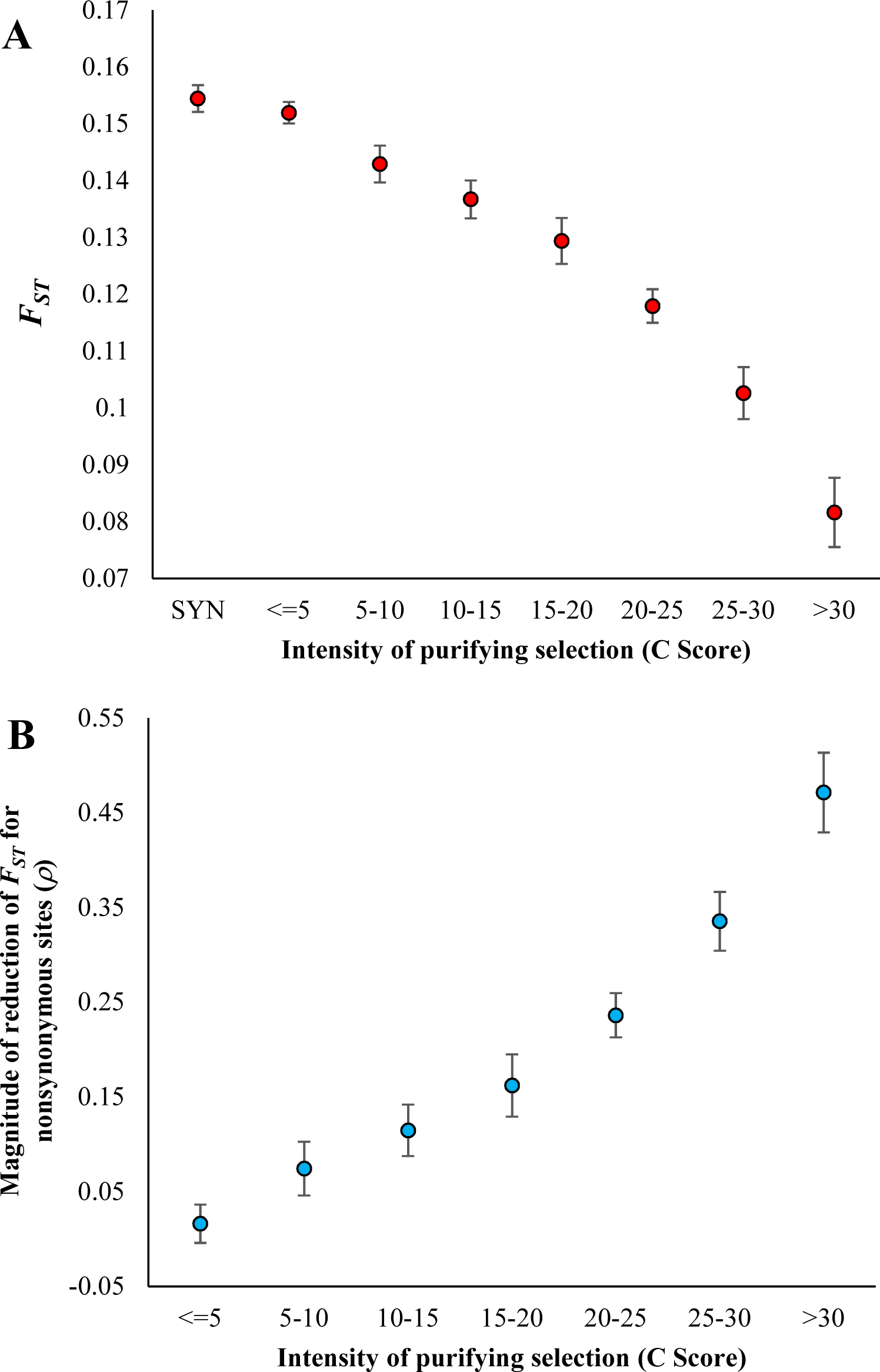
**(A)** Relationship between selection intensity and *F*_ST_. Whole exome data comprising synonymous SNPs (sSNPs) and nonsynonymous SNPs (nSNPs) for the Italian (TSI)-Nigerian (YRI) population pair was used to estimate *F*_ST_. The magnitude of selection intensity on nSNPs is measured by the Combined Annotation-Dependent Depletion (CADD) method that integrates many diverse annotations into a single measure (*C* score) (Kircher, et al. 2014). A bootstrap resampling procedure (1000 replicates) was used to estimate the standard error. **(B)** Magnitude of reduction of *F*_ST_ estimates and selection intensity. X-axis shows the reduction in *F*_ST_ estimates of nSNPs in comparison with that of sSNPs (*ρ*) using equation 11 (see methods) for the exome data described above. Error bars show standard error of the mean.

### Relationship between *F*_ST_ at neutral and constrained genomic regions

To understand the actual cause of the reduction in *F*_ST_ for constrained SNPs we examined the theoretical relationship between *F*_ST_ at neutral and constrained regions. We showed that the fractions of neutral+deleterious variations segregating between (*f*_b_) and within (*f*_w_) populations hold the answer to this. If these fractions were similar (*f*_b_ = *f*_w_) then FST estimates for sSNPs and nSNPs are expected to be equal (Equation 5). However, it is well known that a higher proportion of slightly deleterious SNPs is expected to segregate within populations than that of between populations. This is because a significant fraction of them are purged by purifying selection over time and hence their fraction gets diminished for between population comparisons. Therefore, we show that the *F*_ST_ estimated for nSNPs is expected to be smaller than that of sSNPs as the fraction of neutral+deleterious SNPs segregating within populations is higher than that segregates between populations (*f*_w_ > *f*_b_) (Equation 7). To quantify the magnitude of difference between the two fractions we proposed the measure *η*, which is the excess fraction of deleterious SNPs segregating within populations than that present between populations (Equations 8&9). We show the relationship between *η* and the magnitude of reduction of *F*_ST_ estimated for nSNPs compared to that of sSNPs (*ρ*) (Equation 12), which clearly shows that *ρ* is depended on *η*.

Using the within (*H*_w_) and between (*H*_b_) population heterozygosities for the sSNPs of the European-African comparison, the theoretical relationship between *ρ* and *η* (based on equation 12) was plotted. Figure 2 reveals a positive correlation between the two variables. The values of *ρ* estimated for the nSNPs (using equation 11) belonging to the seven selective constraint categories (*C*-scores) were overlaid on the theoretical line and the corresponding *η* values were predicted (red dots on the line). This suggests that for highly constrained SNPs (*C*-score >30) there is 8.6% excess fraction of deleterious SNPs is present within populations compared to that segregating between populations and this results in a 47% reduction in the *F*_ST_ estimate. This excess was only 0.3% for the SNPs under relaxed selective constraints (*C*- score <=5), which resulted in a 2% reduction in the *F*_ST_. Hence these results suggest that the magnitude of reduction is indeed dictated by the excess fraction of deleterious SNPs segregating within populations compared to that segregate between populations.

**Figure 2.**
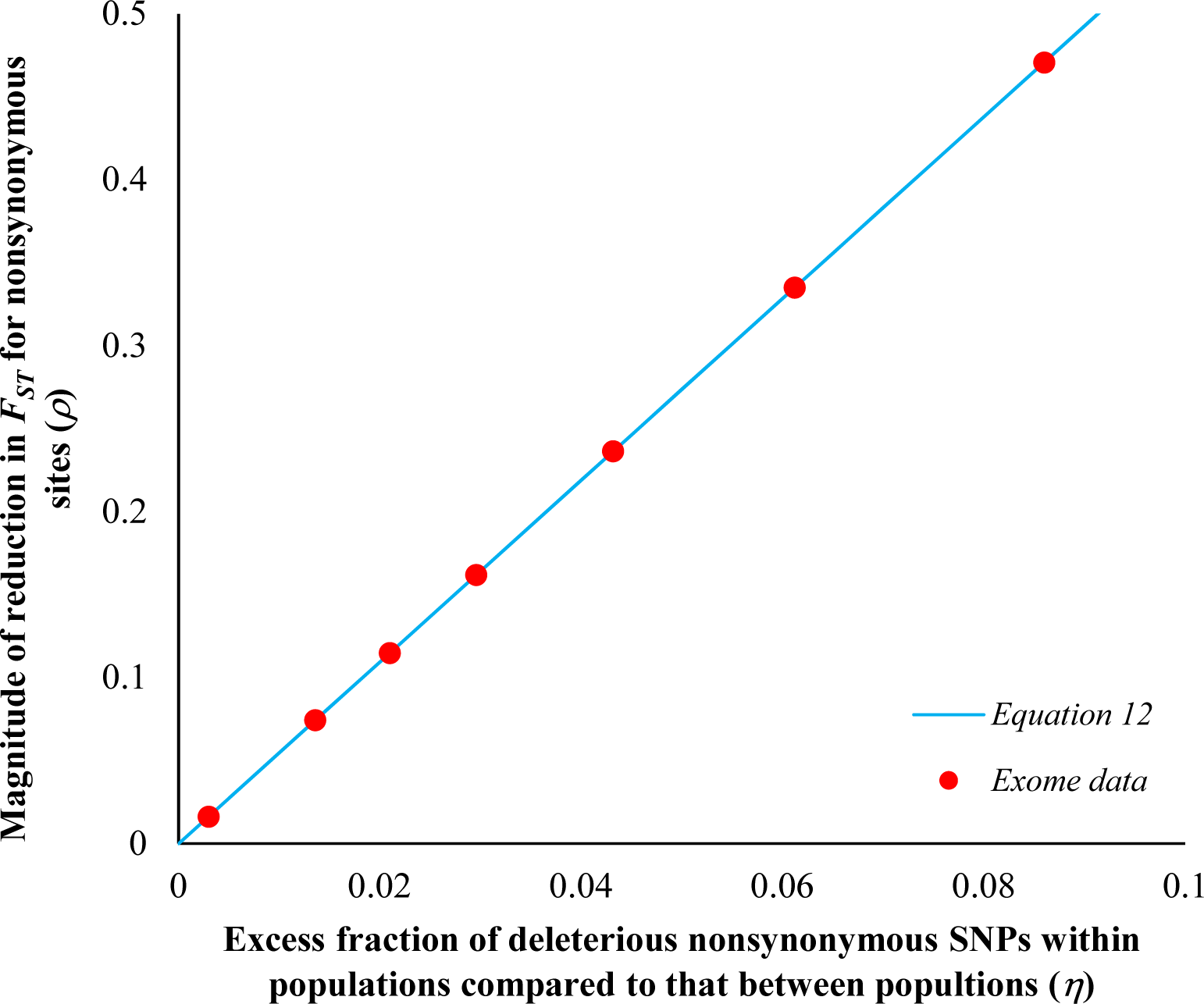
Theoretical relationship between the excess in the fraction of deleterious mutations segregating within population than that of between populations (*η*) and the magnitude of reduction in the *F*_ST_ estimates of nSNPs (*ρ*) using equation 12 (see methods). The line was plotted using within and between population heterozygosities of neutral sSNPs for the Italian-Nigerian comparison and the red dots are the *ρ* values estimated from the exome data using equation 11. Using the theoretical expected line, *η* values were predicted for the corresponding observed *ρ* values.

### *F*_ST_ estimates and population divergence

Next, we investigated the effects of purifying selection on *F*_ST_ with respect to population divergence. This is to compare the magnitude of reduction of *F*_ST_ estimated for closely and distantly related populations. For this purpose, we used four pairs of comparisons with different levels of divergence, European (Italian/TSI) – African (Nigerian/YRI), European (Italian/TSI) – Asian (Chinese/CHB), Southern European (Italian/TSI) – Northern European (British/GBR) and two Southern Europeans (Italian/TSI – Spanish/IBS). Figure 3 (A-D) shows *F*_ST_ estimates obtained for sSNPs and highly constrained nSNPs for the four pairs of populations. This pattern suggests that there is a positive correlation between the population divergence and the extent of reduction of *F*_ST_, which is clear in Figure 4. The *F*_ST_ observed for constrained nSNPs of the distantly related Italian-Nigerian pair was 47% smaller than that of sSNPs (Figure 4). While this reduction was 30% for the Italian-Chinese pair and 16% for Italian-British comparison, it was only 4% for the closely related Italian-Spanish pair (Figure 4). We then examined the theoretical relationship by plotting the relationship between *ρ* and *η* (Equation 12) for the four pairs populations. For this purpose, we used the within and between population heterozygosity estimates of sSNPs of Italian-Nigerian, Italian-Chinese, Italian-British and Italian-Spanish populations. While all four relationships show a positive trend between *ρ* and *η*, there was a huge difference in the slopes of these relationships. The slopes observed for closely related pairs are much higher than that of the distantly related pair. Using the expected theoretical lines, the corresponding *η* values were predicted for the *ρ* values estimated for the four pairs of populations (red dots on the lines). This analysis showed that the excess fractions of 8.6%, 4.0%, 0.2% and 0.03% slightly deleterious nSNPs are present in within populations than between populations of the Italian-Nigerian, Italian-Chinese, Italian-British and Italian-Spanish pairs respectively. The presence of these excess fractions resulted in 47%, 30%, 16% and 4% reduction in the *F*_ST_ estimated for the highly constrained nSNPs (*C*-score > 30) of the corresponding pairs of populations respectively.

**Figure 3.**
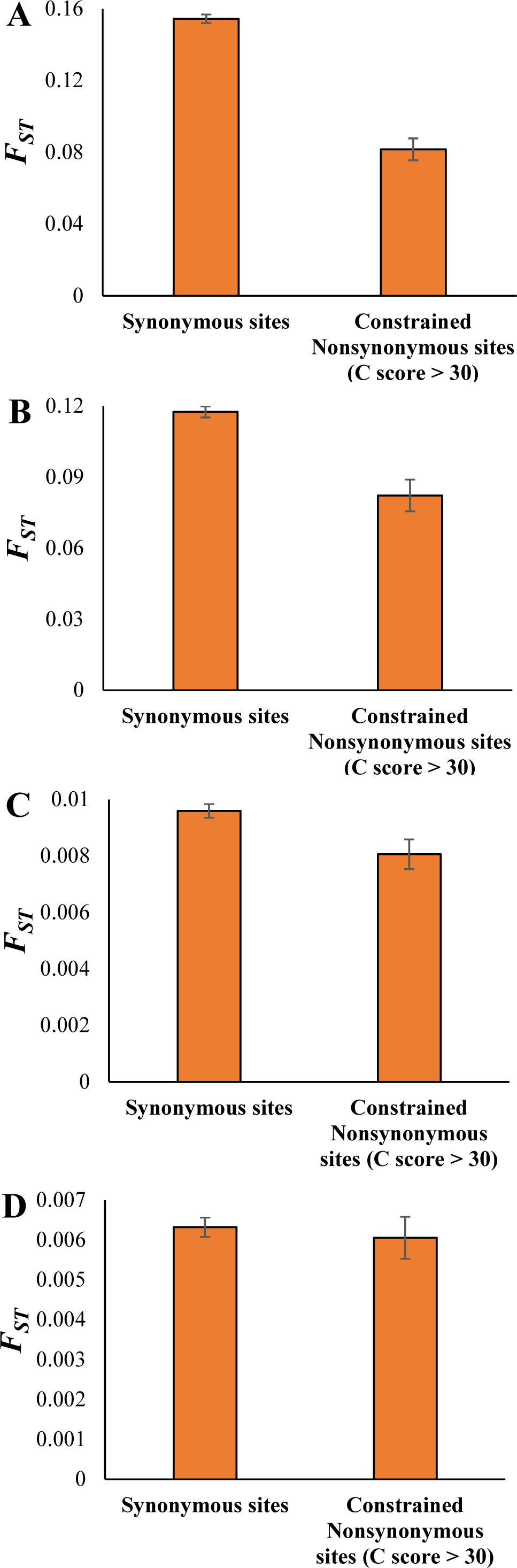
*F*_ST_ estimates for synonymous and highly constrained nonsynonymous SNPs of the **(A)** Italian-Nigerian **(B)** Italian-Chinese **(C)** Italian-British and **(D)** Italian-Spanish population pairs. Error bars are the standard error of the mean and a bootstrap resampling procedure (1000 replicates) was used to estimate the variance. The difference between the FST estimates of neutral and constrained sites are highly significant (P < 0.01, *Z* test) for three comparisons and not significant for the Italian-Spanish pair.

**Figure 4.**
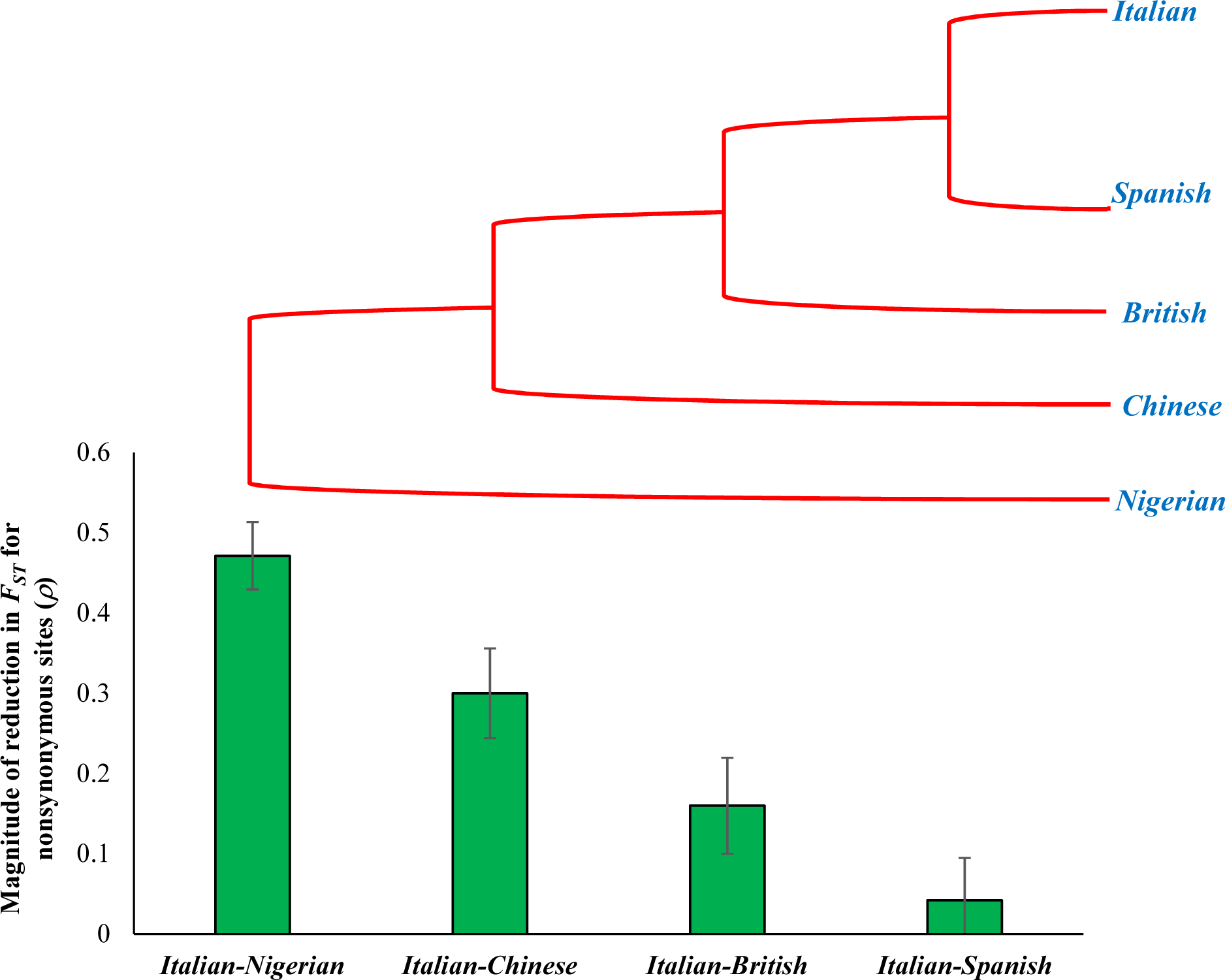
The magnitude of reduction in *F*_ST_ estimates of nSNPs obtained for four population pairs. The population tree on top is drawn to highlight the correlation between the population divergence and the magnitude of reduction in *F*_ST_.

**Figure 5.**
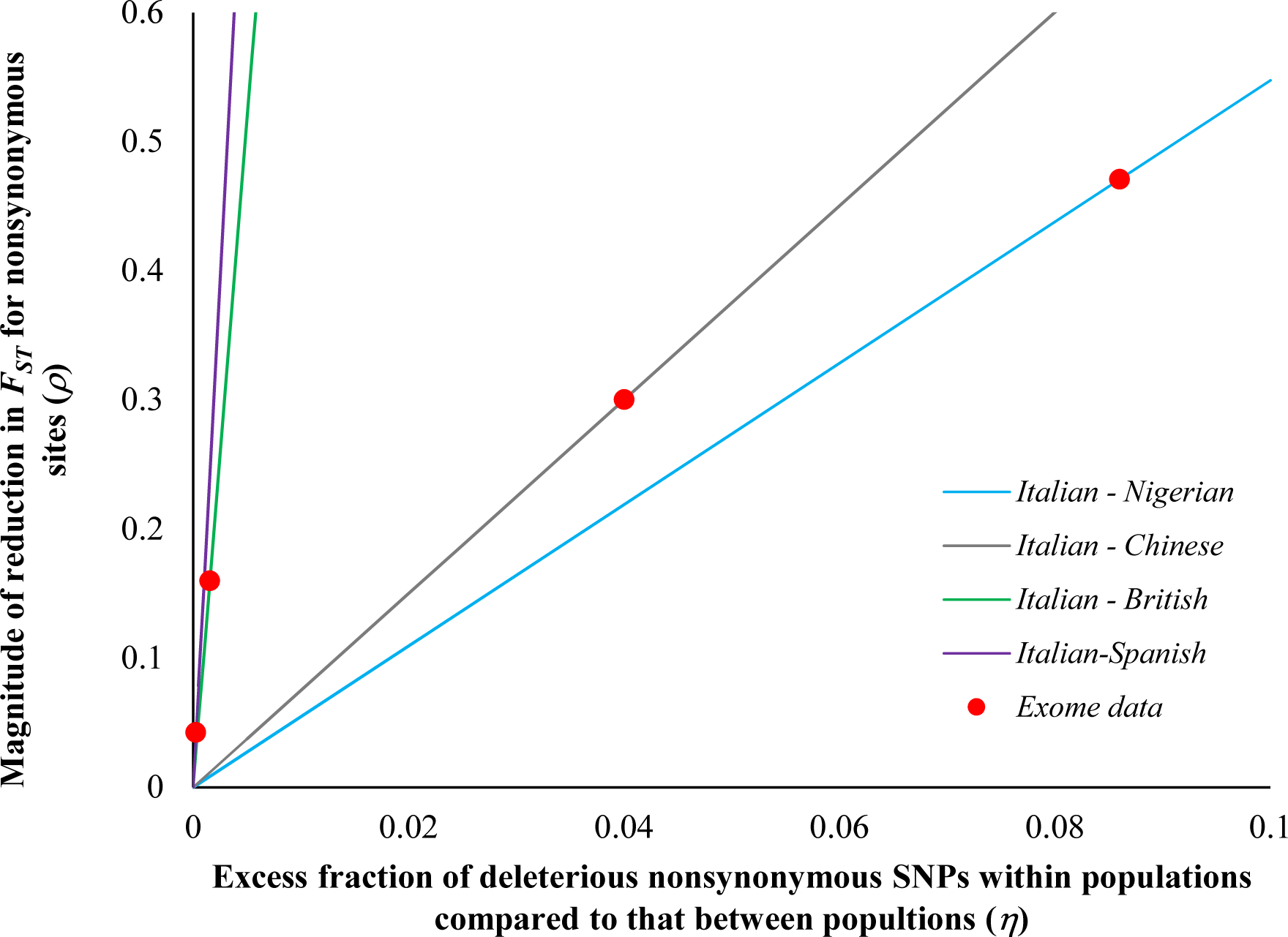
Theoretical relationship between the excess in the fraction of deleterious mutations segregating within population than that of between populations (*η*) and the magnitude of reduction in the *F*_ST_ estimates of nSNPs (*ρ*) using equation 12 (see methods). Neutral population diversities based on sSNPs for within and between population comparisons of the Italian-Nigerian, Italian-Chinese, Italian-British and Italian-Spanish pairs were used to plot the lines and the *ρ* estimated from the exome data (using equation 11) are shown as red closed circles. The theoretical lines were used to predict the *η* values for corresponding *ρ* estimated using the exome data.

## Discussion

Although previous studies have observed a reduction in *F*_ST_ estimates of selectively constrained sites (Barreiro, et al. 2008; Maruki, et al. 2012; Jackson, et al. 2015) the true cause for that reduction was established in this study. Using the theoretical relationship between *F*_ST_ at neutral and constrained sites we showed that an excess fraction of deleterious mutations segregating within population compared to that between populations (*η*) is the reason for the reduction in *F*_ST_ at constrained sites. We also showed the relationship between *η* and the magnitude of reduction in the *F*_ST_ of constrained nSNPs in comparison with that of neutral sSNPs (*ρ*). The reason for the excess fraction *η* present within populations is due to that fact that a high proportion of deleterious mutations segregating within populations are relatively young and hence were not removed by natural selection. Therefore, they contribute significantly to the constrained site heterozygosity within populations. In contrast, a much higher proportion of the harmful mutations have been purged due to the time elapsed and hence their contribution to the constrained site heterozygosity between populations is relatively less. Hence within population heterozygosity at constrained sites is much more inflated than that observed for inter-population comparison. This results in the reduction of *F*_ST_ estimates, as it is based on the normalized difference between the inter- and intra-population diversities.

The results of this study highlight two important patterns and provide theoretical and empirical explanations for them. First the reduction in the *F*_ST_ estimates positively correlates with the magnitude of selection suggesting a much higher underestimation for nSNPs at highly constrained regions of the genome. This is because high magnitude of purifying selection leads to segregation of more slightly deleterious mutations within populations and hence the fraction of deleterious nSNPs segregating within populations will be much higher than that segregates between populations (*f*_w_ ≫ *f*_b_ or *η* ≫ 0). Hence this leads to a much higher underestimation of *F*_ST_ of nSNPs at highly constrained regions compared to that of sSNPs (*F*_ST(N)_ ≪ *F*_ST(S)_). In contrast, there are fewer deleterious nSNPs in less constrained regions and hence the fraction of harmful polymorphisms segregating within populations is expected to be only modestly higher than that segregates between populations (*f*_w_ > *f*_b_ or *η* > 0). This results in a much smaller reduction in the *F*_ST_ estimated for nSNPs present in regions under relaxed selective constraints (*F*_ST(N)_ < *F*_ST(S)_).

Second, we have shown that the magnitude of reduction of *F*_ST_ at constrained sites for comparisons involving distantly related populations was much higher than that of those involving closely related pairs. For instance, this reduction for the European-African comparison (47%) is more than ten-fold higher than that of the Southern European pair (Italian-Spanish) (4.2%) pair. It is well known that deleterious variants are removed over time and hence the only a small fraction (*f*_b_ ≪ 1) of them segregate and contribute to constrained site inter-population diversity for distantly related populations. However, a relatively modest fraction (*f*_b_ < 1) of harmful nSNPs contribute to the inter-population diversity for closely related population as the elapsed time not enough to purge most of them. On the other hand, the fraction of deleterious nSNPs within population (*f*_w_) remain the same for comparisons involving both distantly as well as closely related populations. Therefore, the excess fraction *η* (which is the normalized difference between *f*_w_ and *f*_b_) is much smaller for the comparisons involving closely related populations (*η* ≪1) than those involving distantly related populations (*η* <1). Hence the magnitude of reduction in constrained site *F*_ST_ (with respect to neutral site *F*_ST_) for distantly related populations (eg. European-African) is much higher (*F*_ST(N)_ ≪ *F*_ST(S)_) than that observed for closely related populations (*F*_ST(N)_ < *F*_ST(S)_) (eg. Italian-Spanish).

In this study we used the Hudson et.al formula (Hudson, et al. 1992) to derive the relationship between *F*_ST_ at neutral and constrained sites and also to estimate *F*_ST_ from exome data. This method compares heterozygosities in between and within populations. In contrast Nei (Nei 1973) developed a method that compares heterozygosities of total and subpopulations. Therefore, we derived the relationship between *F*_ST_ at neutral and constrained sites for the method of Nei as well (Equation 13) and repeated all analyses using Nei’s estimator (Supplementary Figures S1-S3). However, this analysis produced similar results to that obtained using the method of Hudson et.al.

The findings of this study suggest that the *F*_ST_ estimated for different genes or genomic regions of a genome are not comparable if the level of selective constrains are different between them. This is particularly important while using *F*_ST_ estimates to detect positive selection because such methods assume neutral evolution in genes and genomic regions and hence do not account for purifying selection in the estimations (Beaumont and Balding 2004; Xue, et al. 2009; Bonhomme, et al. 2010; Chen, et al. 2010; Wu and Zhang 2010; Vitti, et al. 2013). Our results also strongly indicate that *F*_ST_ obtained from the constrained regions of different pairs of populations are not comparable if the population divergence times between the pairs are not the same. In such cases *F*_ST_ estimations should include only neutral sites to obtain unbiased estimates. However, this is only possible for large genomes such as vertebrate in which constrained regions constitute only a small fraction (∼10%) of the genome (Ponting and Hardison 2011; Rands, et al. 2014). This is an important issue for small genomes such as those of fruit flies with >50% of the genome is under selection (Andolfatto 2005). The whole genome based *F*_ST_ obtained for a closely related pair of fruit fly populations is expected to be much smaller than that that obtained for distantly related populations. Therefore, population divergence time need to be considered when comparing the genome-wide *F*_ST_ estimates from different populations. This is particularly more important for invertebrates, protists and prokaryotes as a large proportion of their genomes is under selection.

## Acknowledgments

The author acknowledges the support from the Australian Research Council (LP160100594) and the University of the Sunshine Coast.

