## Supplemental Figure 1-3 for "Temporal distribution of deleterious variations influences the estimation of *F*_ST_"

Sankar Subramanian

*GeneCology Research Centre, The University of the Sunshine Coast, 90 Sippy Downs Drive, Sippy Downs Qld 4556, Australia*

| Intensity of purifying selection (C Score) | FST |
| --- | --- |
| >30 | 0.0425597 |
| 25-30 | 0.0540923 |
| 20-25 | 0.0626702 |
| 15-20 | 0.0691786 |
| 10-15 | 0.0733722 |
| 5-10 | 0.0769721 |
| <=5 | 0.0822251 |
| SYN | 0.0836867 |

| Intensity of purifying selection (C Score) | Magnitude of reduction of FST for nonsynonymous sites (r) |
| --- | --- |
| >30 | 0.491441 |
| 25-30 | 0.353634 |
| 20-25 | 0.251134 |
| 15-20 | 0.173363 |
| 10-15 | 0.123251 |
| 5-10 | 0.0802356 |
| <=5 | 0.0174652 |

**Figure S1. (A)** Relationship between selection intensity and *F*_ST_ using Nei’s estimator (*G*_ST_). Whole exome data comprising synonymous SNPs (sSNPs) and nonsynonymous SNPs (nSNPs) for the Italian (TSI)-Nigerian (YRI) population pair was used to estimate *F*_ST_. The magnitude of selection intensity on nSNPs is measured by the Combined Annotation-Dependent Depletion (CADD) method that integrates many diverse annotations into a single measure (*C* score). **(B)** Magnitude of reduction of *F*_ST_ estimates and selection intensity. X-axis shows the reduction in *F*_ST_ estimates of nSNPs in comparison with that of sSNPs (*ρ*) using equation 11 (see methods) for the exome data described above. Error bars show standard error of the mean.

|  | FST |
| --- | --- |
| Synonymous sites | 0.00481915 |
| Constrained Nonsynonymous sites (C score > 30) | 0.00404516 |

|  | FST |
| --- | --- |
| Synonymous sites | 0.00317098 |
| Constrained Nonsynonymous sites (C score > 30) | 0.00303644 |

|  | FST |
| --- | --- |
| Synonymous sites | 0.0624502 |
| Constrained Nonsynonymous sites (C score > 30) | 0.0429091 |

|  | FST |
| --- | --- |
| Synonymous sites | 0.0836867 |
| Constrained Nonsynonymous sites (C score > 30) | 0.0425597 |

**Figure S2.** *F*_ST_ estimates (using Nei’s estimator, *G*_ST_) for synonymous and highly constrained nonsynonymous SNPs of the **(A)** Italian-Nigerian **(B)** Italian-Chinese **(C)** Italian-British and **(D)** Italian-Spanish population pairs. Error bars are the standard error of the mean. The difference between the FST estimates of neutral and constrained sites are highly significant (P < 0.01, *Z* test) for three comparisons and not significant for the Italian-Spanish pair.

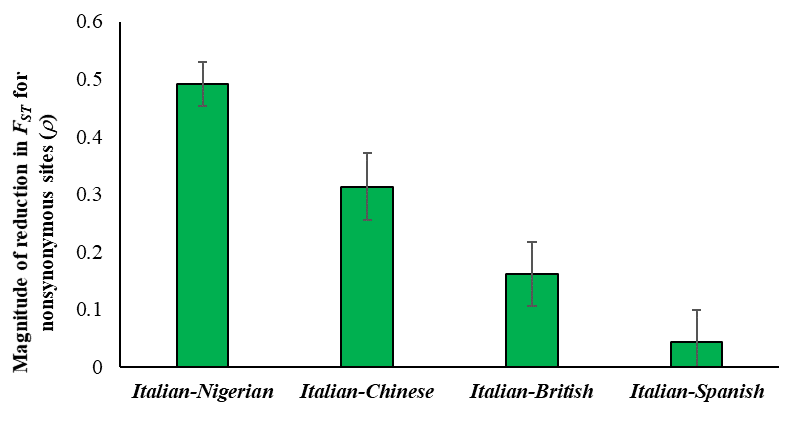

***Italian***

***British***

***Chinese***

***Nigerian***

***Spanish***

|  | Magnitude of reduction in FST for nonsynonymous sites (r) |
| --- | --- |
| Italian-Nigerian | 0.491441 |
| Italian-Chinese | 0.312907 |
| Italian-British | 0.160609 |
| Italian-Spanish | 0.0424269 |

**Figure S3.** The magnitude of reduction in *F*_ST_ estimates (using Nei’s estimator, *G*_ST_) of nSNPs obtained for four population pairs. The population tree on top is drawn to highlight the correlation between the population divergence and the magnitude of reduction in *F*_ST_.
